# Factors behind poor cognitive outcome following a thalamic stroke

**DOI:** 10.1101/2024.07.24.604322

**Authors:** Julie P. C. Vidal, Lola Danet, Germain Arribarat, Jérémie Pariente, Patrice Péran, Jean-François Albucher, Emmanuel J. Barbeau

**Affiliations:** Brain and Cognition Research Center (UMR 5549), CNRS, University of Toulouse, Toulouse, France; Toulouse Neuroimaging Center (UMR 825), Inserm, University of Toulouse, Toulouse, France; Neurology Department, CHU Toulouse Purpan, Toulouse, France

## Abstract

**Objective:** Thalamic strokes produce a range of neurological, cognitive, and behavioral symptoms depending on the thalamic nuclei involved. While thalamic strokes are traditionally associated with severe cognitive deficits, recent studies suggest more modest impairments. This study aims to identify the factors that influence the severity of cognitive impairment following thalamic stroke.

**Methods:** We recruited 40 patients (median age 51) with chronic isolated thalamic stroke and 45 healthy subjects. All subjects underwent neuroimaging and neuropsychological testing. Cluster and principal component analyses were used to discriminate patients from healthy subjects based on cognitive performance. Disconnectome maps and cortical thickness were analyzed to understand the distant impact of thalamic strokes.

**Results:** Two cognitive profiles emerged. Cluster 1 included mostly healthy subjects (n = 43) and patients with no or minor deficits (n = 20); Cluster 2 included patients (n = 19) and 2 healthy subjects with severe deficits of verbal memory, executive functions, and attention. Cluster 1 included all patients with right thalamic stroke. Cluster 2 included all patients with bilateral stroke or mammillothalamic tract disruption. Patients with left-sided stroke were equally divided between Cluster 1 and 2. Other significant differences included age, education, interthalamic adhesion disruption, lesion volume, and location. Disconnectome maps showed larger disruptions of the anterior thalamic projection in patients with left-sided stroke of Cluster 2.

**Interpretation:** Contrary to common expectations, our findings indicate that many patients with thalamic stroke have relatively good cognitive outcomes. In contrast, we identified some of the factors behind poor outcomes that may help clinicians.

## Introduction

Given its high vascularity [1], the thalamus is vulnerable to stroke, giving rise to diverse neurological, cognitive and behavioral manifestations depending on which vascular territory and thalamic nuclei are implicated [2]. Historically, our understanding of thalamic lesions has primarily been based on case studies, and these lesions have been linked to severe cognitive deficits, such as amnesia [3], aphasia [4], spatial hemineglect [5], homonymous lateral hemianopia [6] and auditory agnosia [7], among others.

However, recent publications have reported relatively modest cognitive deficits in patients with thalamic lesions. Liebermann et al. [8] studied a group of 68 patients with an isolated thalamic stroke and demonstrated the absence of cognitive differences with the group of healthy subjects using established clinical self-report questionnaires. In another study, the same authors investigated 19 patients, 58% of whom did not exhibit major executive deficits when z-scores were used [9]. Similarly, during visual anterograde memory tests, either moderate or a lack of memory deficits were observed in 75% of 17 patients with chronic left, right or bilateral thalamic lesions [10]. These findings were corroborated by other studies [11; 12]. More recently, a longitudinal study was carried out on 37 patients with an isolated thalamic stroke from the acute to the chronic phase and revealed that patients recovered, at the group scale, no matter which cognitive domain and vascular territory were affected [13].

The discrepancy between earlier research and these more recent studies may be explained by several factors, including the advent of stroke units, which allow the inclusion of patients with subtler deficits in research programs. The development of neuroimaging, in particular high-field MRI, which helps to identify patients with isolated lesions to the thalamus, also goes some way to explaining this discrepancy; without this method, the exact extent of the lesions may have been unknown or underappreciated in earlier studies. A positive bias may also have played a role with a tendency to report compelling cases rather than patients with more modest deficits in earlier cases. Finally, the thalamus is made of both gray matter and white bundles, such as the mammillothalamic tract (MTT, part of Papez’s circuit, [2]) and the interthalamic adhesion [14]. The lack of investigation into the potential disruption of these bundles in cases of thalamic lesions may hinder the accurate interpretation of associated cognitive deficits.

Nevertheless, these recent group studies question our understanding of the role of the thalamus in cognition. They also suggest that there may be several subgroups of stroke patients, some of whom will recover relatively well after their stroke while others will retain significant cognitive deficits. Our study therefore asks: what determines good or poor cognitive outcome following an isolated stroke in the thalamus?

## 2. Materials and Methods

### 2.1 Participants

We collected data from 45 healthy subjects (ages 23-69, median 48.5, 20 males) and 40 patients (ages 23-75, median 51.1, 25 males) with ischemic left, right or bilateral thalamic lesions (L = 28, R = 6, B = 6). These patients were recruited across two studies in the stroke units of the university hospitals of Toulouse. The first study was approved by the Institutional Review Board “Comité de Protection des Personnes Sud-Ouest et Outre-Mer no. 2-11-0”. It included 20 patients under the age of 80 who had experienced a thalamic stroke. Marginal extra-thalamic lesions were accepted in this study (three patients, [15]). The Fazekas and Schmidt score, which assesses white matter lesions, was equal to 1 except for two patients with a score of 2. The second study was authorized by the “Comité de Protection des Personnes Ile-de-France IV” (Ethics Committee). It included 20 patients under the age of 70 with at least one stroke lesion visually reaching the dorsomedian nucleus and no extra-thalamic damage. For both studies, recruitment criteria were detection of a first symptomatic thalamic infarct regardless of complaint or neurobehavioral report before onset and no previously known neurovascular, inflammatory or neurodegenerative diseases. All patients underwent a clinical examination, a neuropsychological assessment and an MRI scan at least three months after their stroke (median: 501, min: 91, max: 2674 days). All data were acquired after obtaining prior written informed consent from the participants. The etiology of each infarct (when identified) and symptoms at onset (sensorimotor and vertebrobasilar, cognitive or both) were collected from each clinical investigation during the acute phase. Healthy subjects were volunteers included in these two studies with no known significant health issues. Non-inclusion criteria were known neurovascular, inflammatory or neurodegenerative diseases.

### 2.2 MRI Acquisition

For the first study comprising 20 patients and 20 healthy subjects, 3D T1-MPRAGE sequences were acquired on a 3T scanner (Philips Achieva) with the following parameters: 1*1*1 mm voxel size, flip angle = 8°, FOV = 240*240, slice number = 170. For the second study comprising 20 patients and 25 healthy subjects, 3D T1-MPRAGE sequences were acquired on a 3T scanner (Philips Achieva) with the following parameters: 0.9*0.9*1mm voxel size, flip angle = 8°, FOV = 256×256, slice number = 189.

### 2.3 Neuropsychological Assessment

The following tests were used: the Free and Cued Selective Reminding test [16] (verbal anterograde memory); the DMS48 task [17] (visual anterograde recognition memory); the Stroop test [18] (inhibition); literal and semantic fluencies [18] (executive functions); D2 [19] (attention); digit-symbol test [20] (working memory); ExaDé confrontation naming test [21] (language), visuospatial and auditory-verbal digit span [20] (working memory) and three mood and affective scales: the State-Trait Anxiety Inventory [22], Starkstein Apathy Scale [23] and Beck Depression Inventory Scale [24].

### 2.4 Neuropsychological analyses

Group comparisons were carried out using a χ^2^ test for nominal data. Neuropsychological test scores were standardized to z-scores using normative scales. The psycho-affective scales were analyzed using raw data due to a lack of adequate normative scales. To minimize multiple comparisons, one subtest by neuropsychological test was selected and the mean z-score by cognitive function was computed resulting in five cognitive functions: working memory (auditory-verbal and visuospatial digit span), verbal memory (FCSRT Delayed Free Recall), executive function (digit-symbol, Stroop interference minus denomination (I-D) response time), language (literal and semantic fluencies, confrontation naming test) and attention (D2 GZ-F, number of processed items minus errors). This selection was based on literature-driven assumptions. In order to reflect a general cognitive performance, the mean of the previous cognitive functions was computed and referred as “Total”.

#### 2.4.1 Cluster analysis

In order to identify subgroups of patients with similar performances independently of the laterality of infarct, a cluster analysis was performed using the kmeans method and by computing the Euclidean distance [25]. Before running this algorithm in both R and Python, we confirmed that our data set contained clusters by assessing the Hopkins clustering tendency and obtained a value of 0.68 (0: uniformly distributed; 0.5: random; 1: highly clustered data). Because cluster analyses don’t support missing values, one patient with a missing value in the attention function was excluded for this analysis. To identify factors and variables that could better explain the clustering, a logistic regression using the stepwise method on JASP was performed.

#### 2.4.2 Principal component analysis

To further explore how patients and healthy subjects could be differentiated by their cognitive performances, a principal component analysis (PCA) using the five previously described cognitive functions in Rstudio (libraries: FactoMineR, factoextra). One patient with a missing value in the attention function was excluded for this analysis. Results are represented by a plot of each individual projected into the space of the two principal components explaining the maximal amount of variance in the dataset. Individuals are colored by clusters. The percentage of explained variance by principal component and the percentages of the contribution of each variable to those principal components are represented in supplementary figure 1 (fviz_contrib, factoextra package).

#### 2.4.3 rmANOVA and Boxplots

To compare healthy subjects and patients with a left, right and bilateral lesion, a repeated measure ANOVA (rmANOVA) was conducted using the z-scores of the five previously defined cognitive functions. Normality was assessed using Shapiro-Wilk’s test and the homogeneity of variance using Levene’s tests. In the event of a grouping effect, we employed a post-hoc t-test. Finally, we used a Kruskall-Wallis tests followed by Bonferroni-corrected Dunn’s tests to quantify which test was the most discriminative and to study the “Total” cognitive function. significant results were represented as boxplots using z-scores by groups in order to analyze individual performances.

### 2.6 Neuroimaging analysis

#### 2.6.1 Lesion Volumetry and Location

Lesions were manually segmented on the native T1w images by two independent investigators using MRIcron software [26]. Lesions masks were generated using the overlap between the segmentation of both investigators (mean dice = 0.84).

Lesions were localized using HIPS-THOMAS (Histogram-based Polynomial Synthesis-Thalamus Optimized Multi-Atlas Segmentation; [27, 28]) as it allows to segment T1w images in their native space.

To confirm the segmentation results, both the Morel atlas (Krauth et al., 2010) and the FreeSurfer methods were used. If a lesion reached the MTT with a volume greater than 5mm^3^, the MTT was considered to be disrupted using the Morel’s segmentation. To enable comparisons between segmentation methods, nuclei were gathered into nuclear groups using the Morel’s atlas repartition (Anterior: AV, AM, AD, LD; Medial: Hb, MD, MV, Pv, CeM, CL, CM, Pf, sPf; Posterior: LGN, Li, LP, SG, MGN, Po, PuA, PuI, puL, PuM; Lateral: VA, VL, VM, VPI, VPL, VPM) [29, 30]. Finally, the mean percentage of lesion affecting each nuclear group by cluster was computed using the three segmentation methods. As data are not normally distributed, the lesioned volumes by nuclear group for all subjects were compared between methods using a Friedman test (paired data) followed by a Durbin Conover test that had undergone Bonferonni correction for multiple comparisons. HIPS-THOMAS segmentations were used to represent the main results, making the results section easier to read.

#### 2.6.2 Normalization

Native T1w-images and segmented lesions were registered to the MNI152 template using an affine and diffeomorphic deformations computed after a skull stripping via the normalization tool of the BCBtoolkit, based on ANTs (Advanced Normalization Tools) [31]. As spatial normalization can be affected by the presence of lesions by creating a mismatch between the patient brain and the template, for the registration of patient’s T1w-images with bilateral lesions, the lesion mask was used to compute the diffeomorphic transformation in the healthy territory of the brain only [32]. For the normalization of patient’s T1w-images with unilateral lesion, an enantiomorphic transformation was computed by replacing the lesioned area with the signal from the contralateral healthy tissue in order to improve the normalization [33]. The normalized lesions resulting from this method were used to create overlapping lesions of all patients and by clusters on the MNI152 template using MRIcroGL.

#### 2.6.3 Disconnectome

To map the impact of thalamic strokes on white matter tracts, we used the Disconnectome maps software, which is part of the BCBtoolkit [34]. Each patient’s normalized lesion was registered onto the diffusion-weighted imaging datasets from a group of ten healthy individuals constituting a reference multi-atlas. These standardized lesions served as seeds in the fiber tracking analyses to identify white matter fibers intersecting with the stroke-affected areas, in comparison with the multi-atlas of white matter tracts. The software output is, for each patient, a map associated with voxel-wise probabilities of disconnection within the standard MNI152 space. It indicates the likelihood of white matter tract disruption. The probability that each tract might be disconnected was given by the Tractotron software from the same toolkit. Tracts with a disconnection probability above 50% were considered to be significantly disrupted [34]. Disconnectome maps were then thresholded at this 50% probability mark to focus on the most relevant disconnections. By averaging these thresholded maps across patients by clusters, we created an average disconnectome map, which was then projected onto an MNI152 template to identify commonly affected tracts.

#### 2.6.4 Cortical thickness

The T1w images were processed using the FreeSurfer image analysis pipeline (http://surfer.nmr.mgh.harvard.edu) to obtain an automatic surface parcellation based on the Desikan-Killiany atlas and compute the cortical thickness for each subject. The mean cortical thickness of the whole brain and by cortical area were extracted from the statistical output file. To answer our hypothesis, Desikan-Killiany ROI were gathered for the whole brain and for specific cortical areas found to be connected to the most disrupted tract identified by the previous analyses. The mean cortical thickness was compared between groups using a Mann-Whitney U test.

## 3. Results

### 3.1 Sociodemographic

Data from 45 healthy subjects and 40 patients who had experienced a focal ischemic thalamic stroke were collected. Groups were comparable in terms of gender (χ^2^ = 2.78, p = 0.10), age at neuropsychological assessment (t-test, p = 0.41) and years of education (t-test, p = 0.12) (Tab.1). The median age at stroke was 51 with a minimum of 21 and a maximum of 75 (Fig. 1A) in our sample. The origin of the thalamic stroke was in almost half of the cases (46%) a congenital anomaly of the atrial septum (Fig. 1B). Initial symptoms at onset were equally divided between motor /vertebrobasilar (28%), cognitive (28%) and both (45%) (Multinomial test: χ^2^ = 2.45; p = 0.3) (Fig.1C). However, right thalamic strokes did not lead to cognitive symptoms at the acute phase, contrary to left or bilateral strokes (Fig. 1D) (χ^2^ = 11.5; p = 0.02).

**Figure 1:**
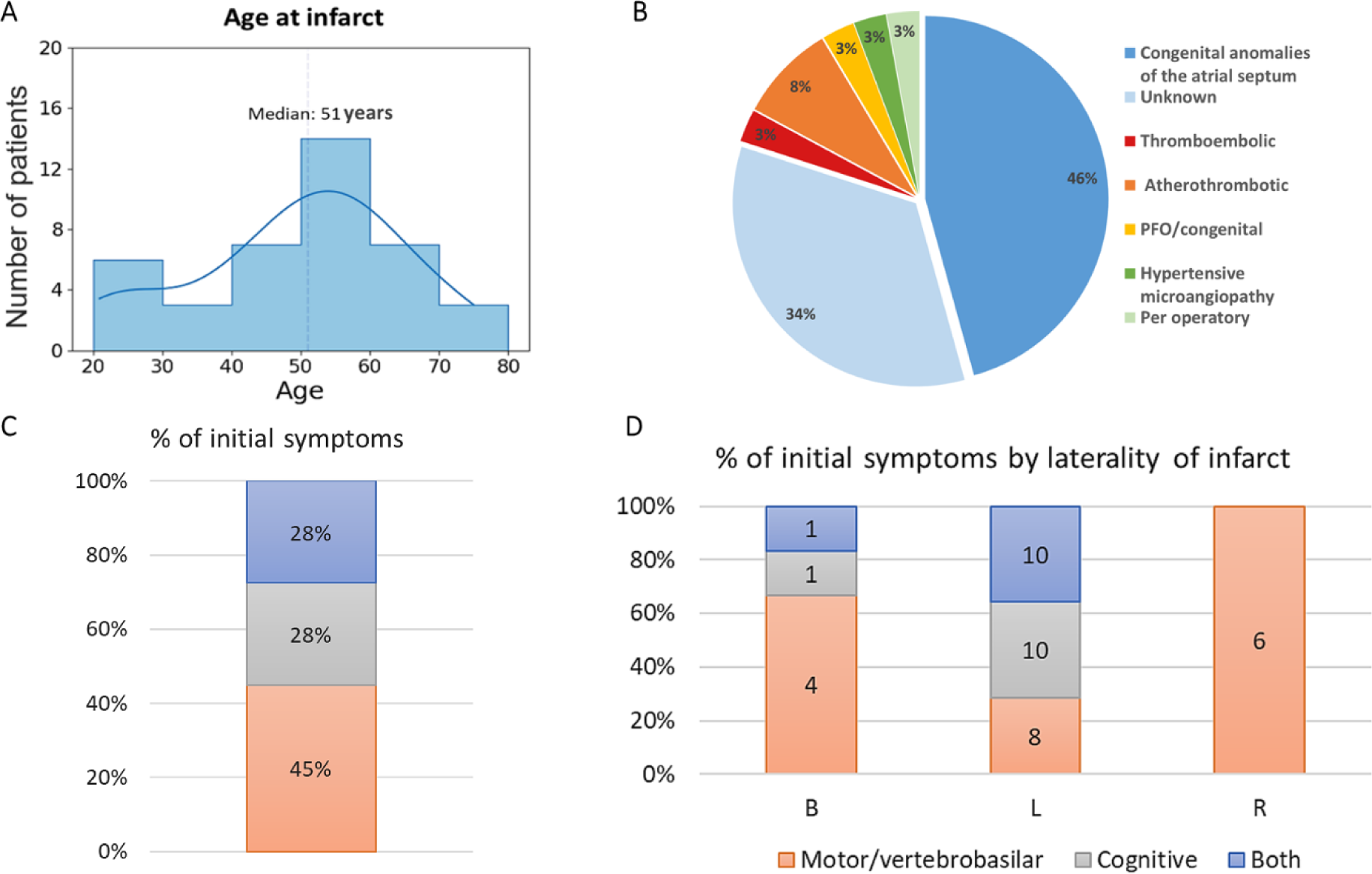
A) Distribution of age at infarct of the patients with an isolated thalamic stroke. B) Percentages of each thalamic stroke etiology. Congenital anomalies of the atrial septum include: atrial septal defect ostium secondum; Patent Foramen Oval (PFO); and/or atrial septal aneurysm. This information could not be found for five patients. This graph therefore concerns 35 patients out of the total of 40. C) Percentages of patients by categories of initial symptoms when the stroke occurred. D) Comparisons of each symptom’s prevalence by laterality of infarct. B: Bilateral; L: Left; R: Right.

### 3.2 Neuropsychological analyses

To identify subgroups of patients and compare them with healthy control subjects, an analysis including all patients and control subjects was conducted using the k-means method on verbal memory, attention, working memory, executive functions and language functions. A two-cluster solution was an optimal number of clusters for portioning the data, obtaining 13/30 indices from the NbClust R package (the second-best option was three clusters, giving 6/30 indices). Cluster 1 included 96% of the healthy subjects and 51% of the patients (n = 20), while Cluster 2 comprised the remaining patients (n = 19) and two healthy subjects (χ^2^ = 22; p < 0.001, comparing the repartition of healthy subjects and patients among clusters).

Patients from Cluster 1 showed no or minor neuropsychological deficits. By definition, these patients were not distinguishable from healthy subjects as they were clustered together. In addition, the median z-scores of neuropsychological performances for people in Cluster 1 were around 0 (Fig. 2A).

**Figure 2:**
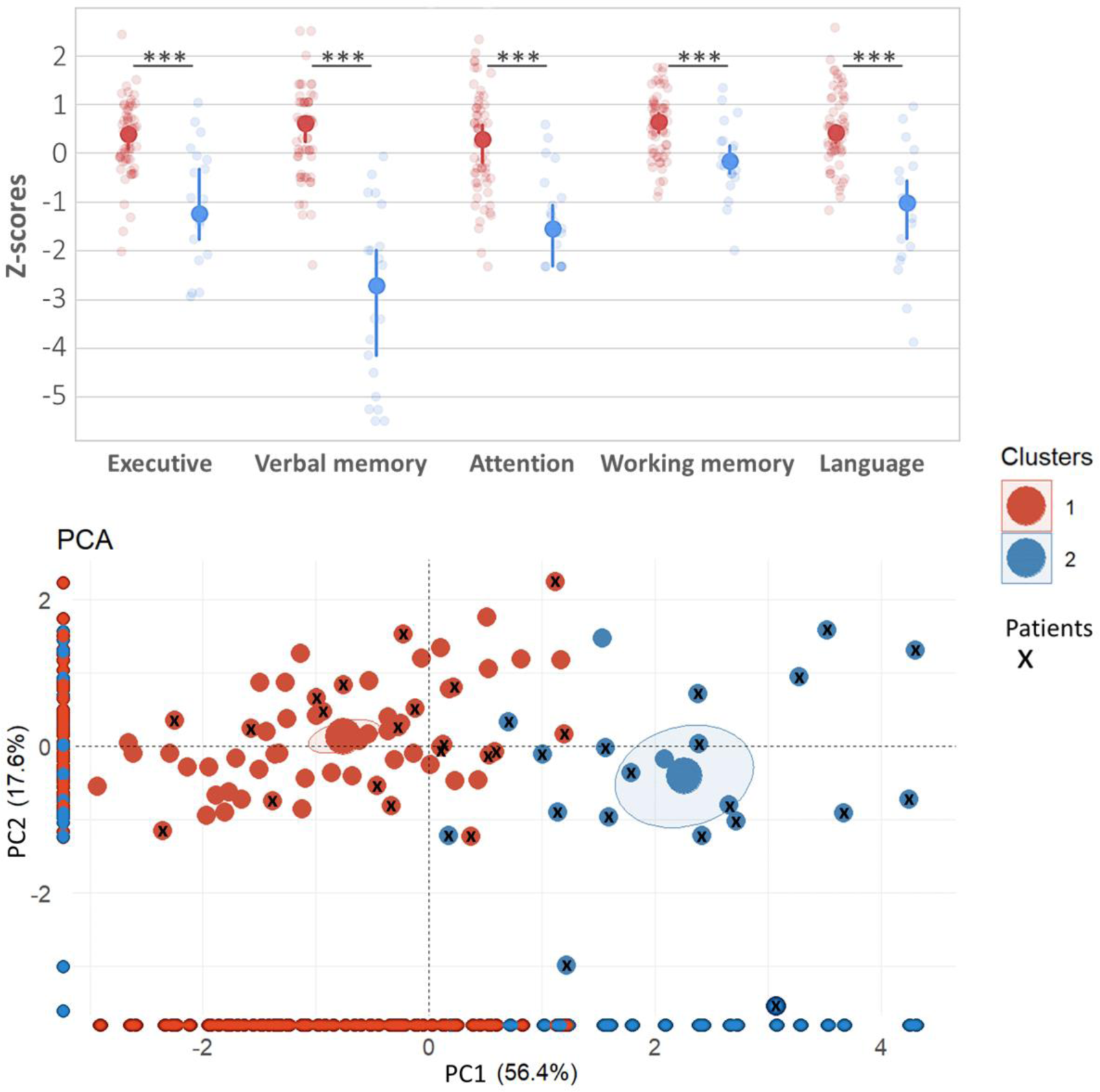
A) Comparisons of the z-scores from the neuropsychological tests between subjects (patients and healthy subjects) of both clusters. For each plot, the solid dot depicts the median while the bar represents the 95% confidence interval. Each dot represents the performance of a subject. The two clusters are significantly different for all cognitive functions (Mann-Whitney, p <0.001). B) Plot of the principal component analysis (PCA). The two larger dots represent the mean by cluster along with their surrounding 95% confidence interval. All subjects are color-coded by cluster from the cluster analysis in panel A. Dots labeled with an “x” represent patients. This labeling is only shown in panel B for better readability. Each individual is projected on the x or y axes to show the ability of each component to distinguish Cluster 1 from Cluster 2.

Several factors differed between both clusters (Tab. 2). Individuals in Cluster 2 were older (albeit young compared to the usual age of stroke patients, 55.6 vs 47.6 years-old, Mann-Whitney, p = 0.025), with a lower number of years of education (Mann-Whitney p = 0.002) and a greater volume of lesions (Mann-Whitney, p < 0.001, mean ± sd (mm^3^): Cluster 1 = 229 ± 214; Cluster 2 = 553 ± 289). The laterality of infarcts was also significantly different between clusters (χ^2^ = 12; p = 0.002). Cluster 1 contained all patients with a right lesion while Cluster 2 contained all patients with bilateral lesions. Likewise, subjects were not equally distributed depending on the presence of, absence of or damage the interthalamic adhesion (IA) (χ^2^ = 8.4; p = 0.015). In Cluster 1, 76% of subjects had an IA and, among them, patients with damage extending into it represented 10% of the dataset, while in Cluster 2, 67% had an IA and patients with a damaged IA represented 43% of the sample. Patients with left lesions were equally distributed among both clusters. Importantly, Cluster 2 contained 100% of the patients with an interrupted MTT. The time between the infarct and both the neuroimaging and the neuropsychological assessment did not differ between both clusters (T.-Test: p-value = 0.74). The etiologies of the infarct (Fig. 1B) did not differ between the patients in each cluster (χ^2^ = 8.3; p = 0.22).

**Table 1:** Sociodemographic information from the dataset.

|  | Healthy subjects | Patients |
| --- | --- | --- |
| **N** | 45 | 40 |
| **Gender (F)** | 25 | 15 |
| **Lesion side (left-right-bilateral)** | - | 28-6-6 |
| **Mean age ± SD (min-max)** | 48.5 ± 13.7 (20-69) | 51.1 ± 14.9 (23-75) |
| **Mean years of education ± SD (min-max)** | 13.58 ± 3.5 (5-21) | 12.4 ± 3.4 (4-17) |

**Table 2:** Comparison of subjects from Cluster 1 and Cluster 2. The lesion volume was computed in the MNI space. Two healthy subjects could not be assessed for the interthalamic adhesion because the two thalami were adhering to each other.

|  | *Cluster 1* | *Cluster 2* |
| --- | --- | --- |
| ***N total subjects /healthy subjects/patients*** | 63-43-20 | 21-2-19 |
| ***Gender (F)*** | 34 | 6 |
| ***Mean age ± sd*** | 47.6 ± 13.9 | 55.6 ± 14.1 |
| ***Mean years of study ± sd*** | 13.7 ± 3.2 | 10.8 ± 3.4 |
| ***Laterality Left-Right-Bilateral*** | 14-6-0 | 13-0-6 |
| ***Interthalamic adhesion (Yes/No/Damaged)*** | 43-13-5 | 8-7-6 |
| ***Mamillothalamic tract lesion*** | 0 | 9 |
| ***Mean lesion volume (mm)<sup>3</sup> ± sd*** | 229 ± 214 | 553 ± 289 |
| ***Days since infarct ± sd*** | 583 ± 630 | 650 ± 612 |

To further explore how patients and healthy subjects could be differentiated by their cognitive performances and identify more discriminative cognitive functions, a PCA was performed on all cognitive functions (attention, executive functions, verbal memory, language). 56.4% of the variance was explained by the first principal component (PC) and 17.6% by the second. The contribution of each variable was almost identical in PC1, while in PC2 the main contribution came from language (42%) and working memory (34%) (Supp. Fig. 1). The resulting projection of each individual’s performances and cluster attribution as presented before into this two-dimensional space is represented in Figure 2B. PC1 allowed us to almost exactly distinguish Cluster 1 from Cluster 2. It also suggests that attention, executive functions and verbal memory are slightly more discriminative and impaired in patients from Cluster 2. This analysis further confirmed the absence or only minor deficits in patients from Cluster 1 (marked as “x” on red dots, Fig. 2B). Only two patients were slightly closer to subjects from Cluster 2 on PC1, but they were also close to the two healthy subjects in this group (absence of “x” on blue dots, Fig. 2B).

A logistic regression analysis was conducted on patients’ data, with cluster as the dependent variable. The aim was to determine which factors from Table 2 best predicted the clustering outcome. This analysis revealed that the most predictive factor for inclusion in Cluster 2 was the interruption of the MTT (χ^2^ = 16, p < 0.001, odds ratio = 4.3*10^-9^). The model achieved an accuracy of 0.95 and an area under the curve of 0.98. Additionally, the second-best predictive model also included the infarct laterality (χ^2^ = 5, p < 0.001). Subsequently, when incorporating the five preceding cognitive functions into the model, the best model was observed when considering only verbal memory (χ^2^ = 36, p < 0.001) and exhibited perfect accuracy and specificity in predicting the clustering outcome. The second-best model also incorporated attention (χ^2^ = 19, p=0.002).

As the laterality of infarct seems to be a determinant factor of the clustering, a rmANOVA was performed on the five cognitive functions explored with the laterality of the stroke as the grouping factor. This resulted in a main effect laterality of infarct (p < 0.001, ηp2 = 0.47). Post-hoc comparisons demonstrated that patients with a bilateral or left stroke were significantly more cognitively impaired than not only healthy subjects (p < 0.001), but also patients with a right thalamic stroke (p < 0.001). Results for each cognitive function are presented in Fig. 3, with individual boxplots presented separately in Supplementary Fig. 2.

**Figure 3:**
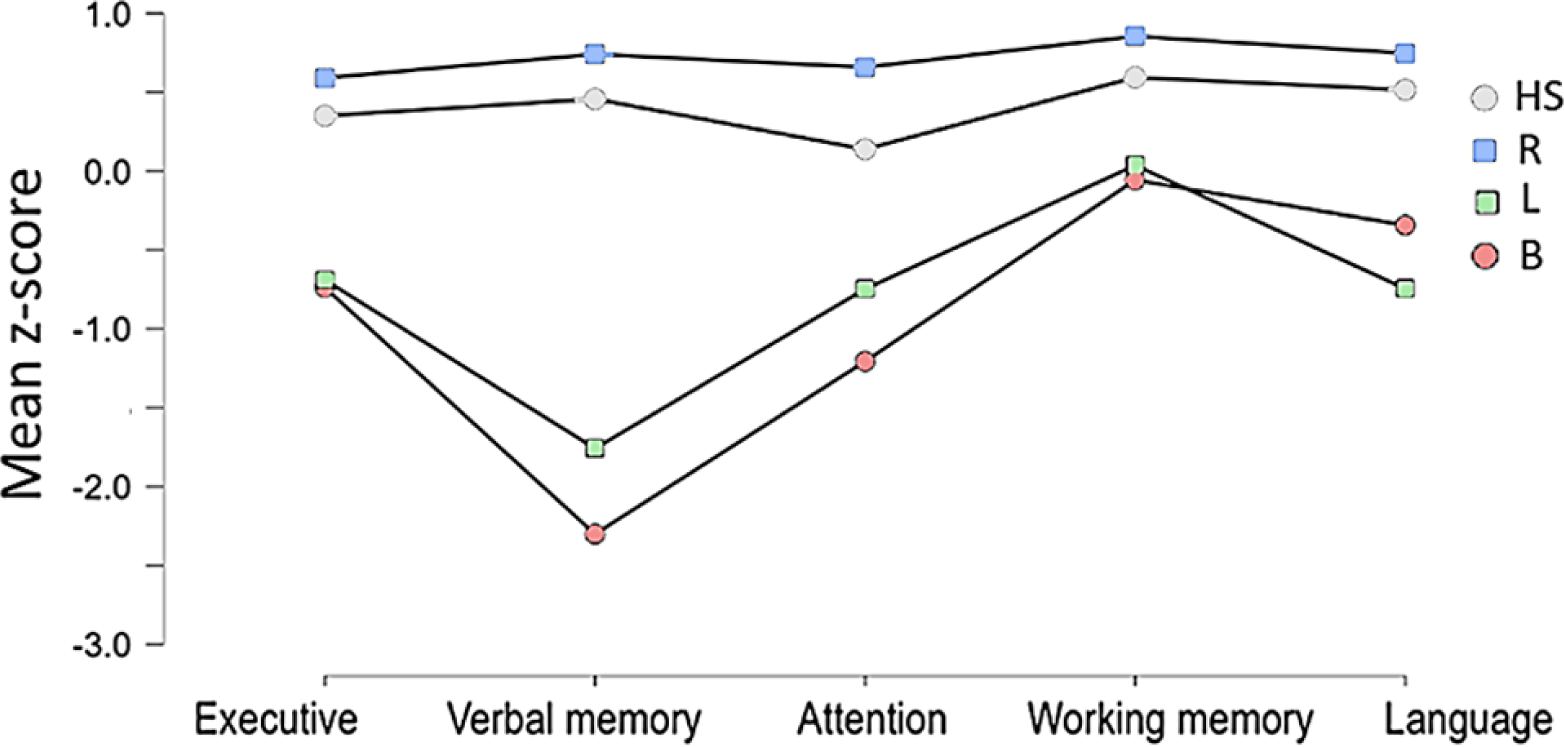
Mean z-score in each cognitive function by groups of patients depending on the laterality of their infarct.

No difference on the psychoaffective scales (Beck, Starkstein, Spielberg; Mann-Whitney, p > 0.3) was demonstrated between healthy subjects and patients considering the laterality of their infarct. When comparing subjects from each cluster, the Starkstein apathy score was significantly higher in subjects from Cluster 2 (Mann-Whitney, p = 0.003). This difference remained significant when comparing only patients from Clusters 1 and 2 (p = 0.01; mean ± SD for Cluster 1: 7.8 ± 4.5; mean ± SD for Cluster 2: 11.4 ± 4.2).

### 3.3 Neuroimaging analyses

A puzzling question related to patients with a left thalamic infarct, as these patients were equally divided between Cluster 1 (n = 14) and Cluster 2 (n = 13). How could the same infarct laterality have no or minor impact on the cognition of some patients and a larger impact in others? We carried out different neuroimaging analyses to tackle this issue. Overlapping lesions by specific clusters and by laterality of infarcts are graphically depicted on the MNI152 template (Fig. 4A). The majority of these lesions were found within the median and lateral left thalamic territories (Fig. 4B). Among patients with left infarcts from Cluster 1, 75% of the volume affected the median left thalamus, while 25% affected its lateral part. In contrast, patients with left infarcts from Cluster 2 showed the opposite pattern (17% vs 83%).

**Figure 4:**
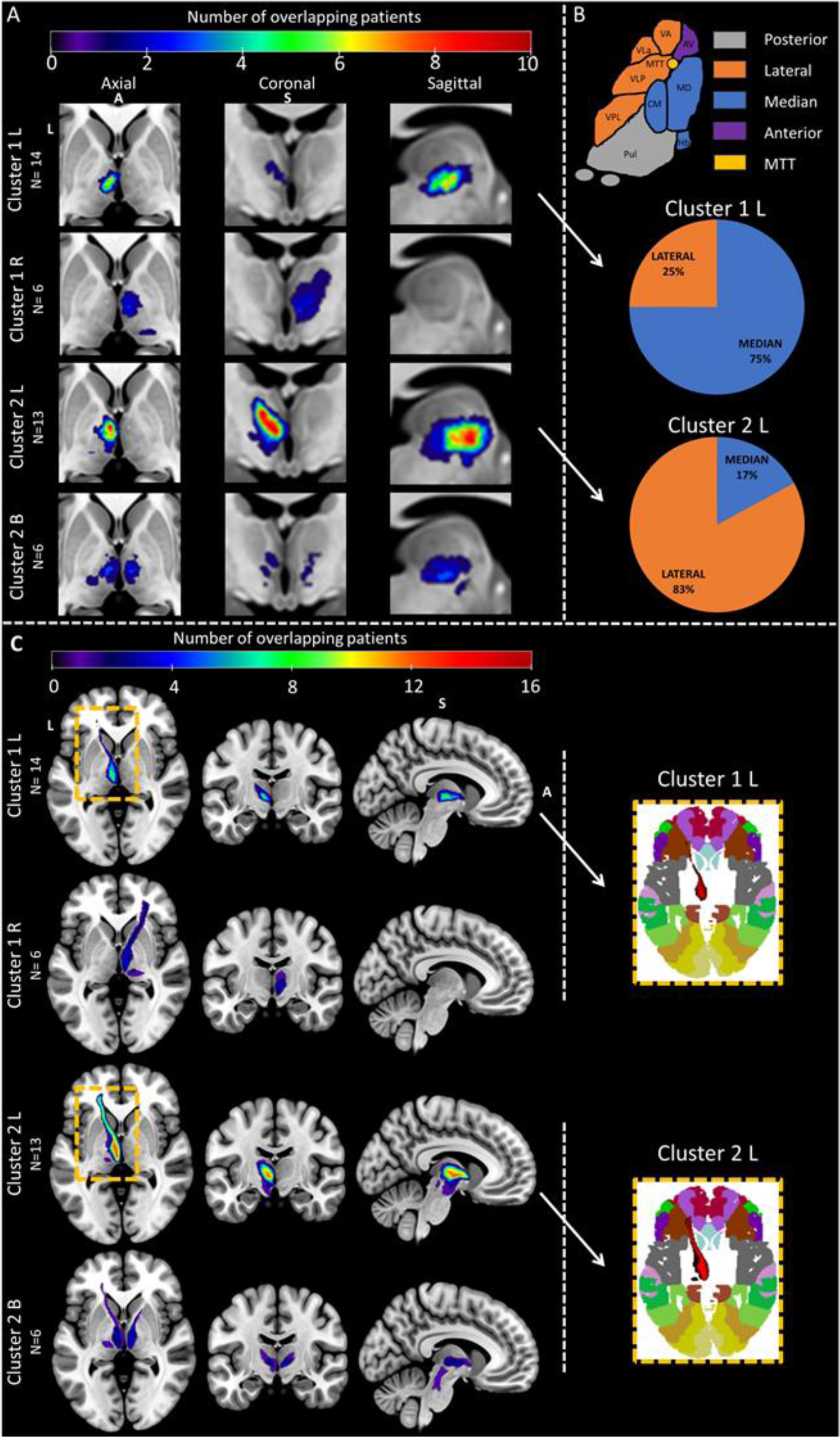
A) Overlapping normalized lesions represented on the same slice of the MNI152 template, which contained the most overlapping lesioned voxels, of patients from Cluster 1 with a left lesion (n = 14) or right lesion (n = 6) as well as patients from Cluster 2 with a left lesion (n = 13) or bilateral lesion (n = 6). Among patients with left infarcts from Cluster 1, 75% of the volume affected the median left thalamus, while 25% affected its lateral part. In contrast, patients with left infarcts from Cluster 2 showed the opposite pattern (17% vs 83%). B) Nuclei repartition into nuclear group (upper figure) and % of lesioned nuclear group of patients with a left lesion from Clusters 1 and 2 using HIPS-THOMAS for the segmentation (bottom figure). C) Mean disconnectome maps by Cluster and by laterality of infarcts with the probability of having an interrupted voxel higher than 50% along with the projection by Cluster on the Brodmann atlas in the MNI152 space (right inset). The disconnection shared by most patients concerned the left anterior thalamic radiations. Patients with a left infarct from Cluster 2 had a more pronounced left anterior thalamic disconnection (right inset in Fig. 4C). When overlapped to the Broadmann atlas, those interrupted radiations projected to areas 11 and 47 respectively, corresponding to the orbitofrontal cortex and the inferior frontal gyrus. A: anterior; S: Superior; L: left. The color bars represent the number of patients with the same lesioned voxels.

To identify not only the local effects of lesions but also their distant effects, a disconnectome map was generated for each patient [34]. This map was thresholded at a 50% probability to isolate voxels representing tracts with a high likelihood of being disrupted. An average disconnectome was then generated for each cluster by laterality of infarct; these are reported in Figure 4C. The disconnection shared by most patients (n = 30, n = 17/19 from Cluster 2 and n = 13/20 from Cluster 1) concerned the left anterior thalamic radiations. The opposite radiation was disconnected in 11 patients (five from Cluster 1 and six from Cluster 2). Patients with a left infarct from Cluster 2 had a more pronounced left anterior thalamic disconnection (right inset in Fig. 4C). When overlapped to the Broadmann atlas, those interrupted radiations projected to areas 11 and 47, corresponding to the orbitofrontal cortex and the inferior frontal gyrus.

As lesions were associated with a high probability of bilaterally interrupting the anterior thalamic projections to the orbitofrontal and ventrolateral prefrontal cortex (vlPFC), the mean cortical thickness of these regions was assessed in both right and left hemispheres along with the anterior cingulate gyrus and the dorsolateral prefrontal cortex (dlPFC), as well as for the whole brain. Indeed, a number of studies have identified that all of these regions are connected with the anterior and mediodorsal nuclei of the thalamus through the anterior thalamic radiations [35, 36]. Note that this analysis included the control subjects as there were too few patients to carry out this analysis in the patient group only. When comparing clusters, subjects from Cluster 2 had significantly inferior mean cortical thickness in the whole brain, as well as in each cortical area of interest (p < 0.005) (Tab. 3).

**Table 3:**
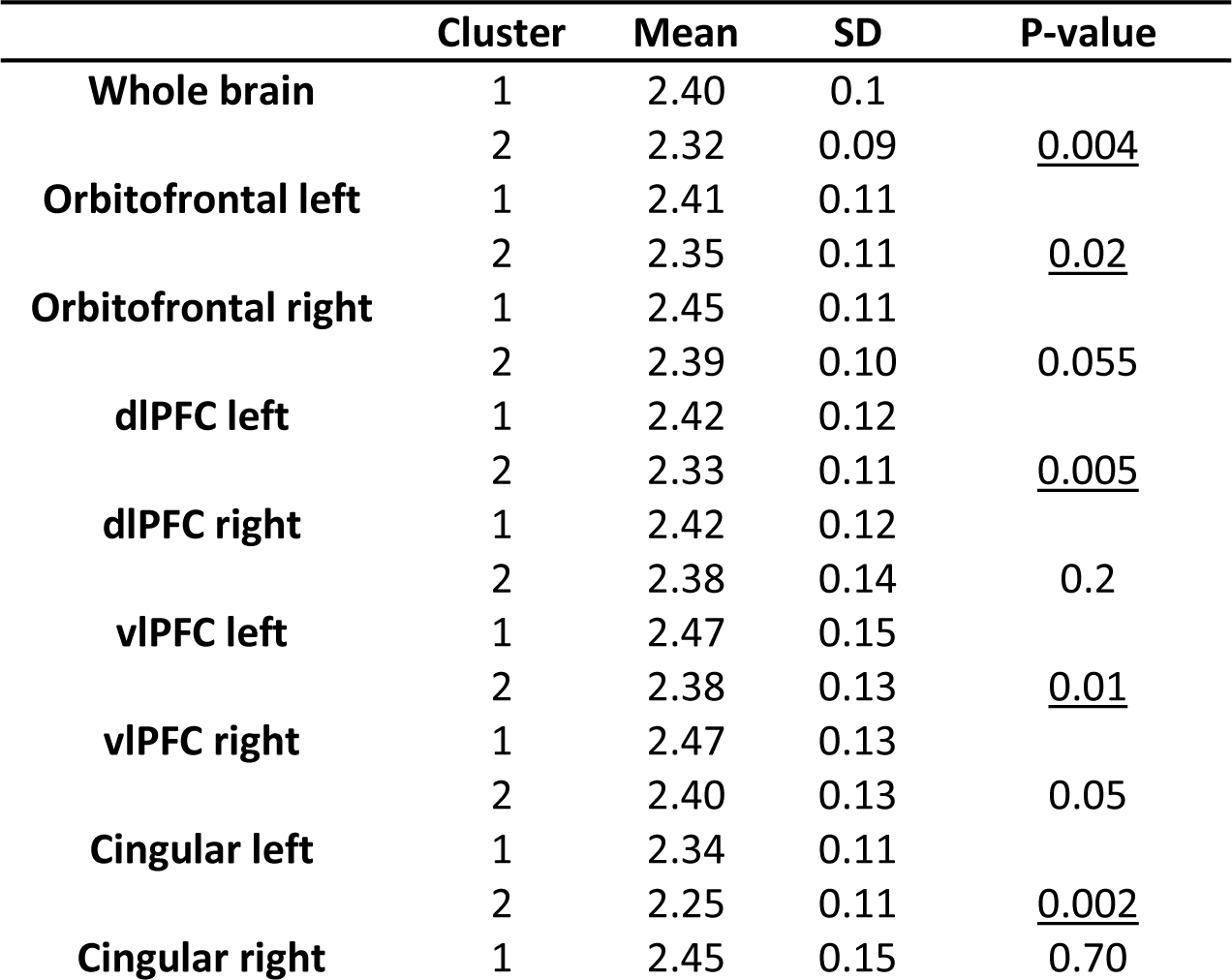
Mean cortical thickness for the whole brain and by cortical parcellations between Clusters 1 (n = 63) and 2 (n = 21) using a Mann-Whitney U test. Significant p-values are underlined.

|  | Cluster | Mean | SD | P-value |
| --- | --- | --- | --- | --- |
| **Whole brain** | 1 | 2.40 | 0.1 |  |
|  | 2 | 2.32 | 0.09 | <u>0.004</u> |
| **Orbitofrontal left** | 1 | 2.41 | 0.11 |  |
|  | 2 | 2.35 | 0.11 | <u>0.02</u> |
| **Orbitofrontal right** | 1 | 2.45 | 0.11 |  |
|  | 2 | 2.39 | 0.10 | 0.055 |
| **dIPFC left** | 1 | 2.42 | 0.12 |  |
|  | 2 | 2.33 | 0.11 | <u>0.005</u> |
| **dIPFC right** | 1 | 2.42 | 0.12 |  |
|  | 2 | 2.38 | 0.14 | 0.2 |
| **vIPFC left** | 1 | 2.47 | 0.15 |  |
|  | 2 | 2.38 | 0.13 | <u>0.01</u> |
| **vIPFC right** | 1 | 2.47 | 0.13 |  |
|  | 2 | 2.40 | 0.13 | 0.05 |
| **Cingular left** | 1 | 2.34 | 0.11 |  |
|  | 2 | 2.25 | 0.11 | <u>0.002</u> |
| **Cingular right** | 1 | 2.45 | 0.15 | 0.70 |

## 4. Discussion

This study set out to explore the neuropsychological outcomes of ischemic thalamic strokes, with a particular focus on identifying some of the factors behind poor cognitive recovery. Our results suggest that cognitive impairments following thalamic strokes may not be as uniform or as severe as previously thought. Our cluster analysis revealed that one subgroup of patients (Cluster 1) showed no or marginal cognitive impairment, in line with what had been suggested in previous studies [8, 9, 10, 11, 12, 13]. This subgroup included all patients with a right thalamus lesion as well as about half of those patients with a left thalamus infarct.

The other subgroup of patients (Cluster 2) showed significant cognitive impairments across several domains, notably in verbal memory, attention and executive functions. These patients also had a higher apathy score. This subgroup predominantly consisted of patients with damage to the left thalamus (49% of patients with a left infarct) as well as all patients with bilateral lesions. This is consistent with the findings of previous studies analyzing cognitive deficits depending on the laterality of infarct [13, 37], as patients with left or bilateral thalamic strokes exhibited more severe cognitive impairments compared to those with right-sided lesions. However, a novelty of our study lies in our highlighting that following left thalamic infarct, about half of all patients will recover well, while the other half will exhibit cognitive deficits.

A major aim of the current study was to analyze the factors that could explain poor outcomes following a thalamus stroke. In addition to the laterality effect, subjects from Cluster 2 were older and had a lower level of education. It is worth noting that the thalamic stroke patients in our dataset are relatively young (56 years old for Cluster 2) compared to the usual stroke population. Patients in Cluster 2 also had larger lesion volume. An interruption of the MTT also systematically led to more severe neuropsychological outcomes. The absence of an the interthalamic adhesion or lesions extending into it was also associated with poorer outcomes.

The intricate relationship between lesion location within the thalamus and resultant cognitive deficits underpins the complexity of thalamic contributions to cognitive functions. Our neuroimaging analysis revealed that lesions predominantly affecting the left thalamus, especially those involving the lateral regions, were associated with more severe cognitive impairments (Cluster 2). A similar anterior left region was also described to lead to mnesic deficits [2, 12, 38] and was associated with severe deficits in language, and executive functions following stroke [13]. A close region was identified by Hwang et al. ([37] Fig. 5A and 8), which led to more pronounced and widespread deficits than patients with lesions elsewhere. Hwang et al. [37] proposed that there is a distinction between connector hubs and provincial hubs within the thalamus, thereby providing a framework that may help us to understand these outcomes. Connector hubs are characterized by their extensive connectivity with multiple cortical networks. To support this analysis, our disconnectome maps revealed that anterior thalamic projections were more extensively disrupted in patients from Cluster 2, particularly when comparing those with left-sided strokes from Cluster 1. Our data thus support the idea that strokes impairing these hub regions, notably in the left lateral thalamus, precipitate widespread cognitive deficits, encompassing verbal memory, attention and executive functions.

In addition, the MTT has been identified as a small but pivotal track for memory impairment [2, 12] that also is located in this same region. Altogether, the relatively small size of this critical region compared to the whole thalamus implies that it has a lower likelihood of being affected; this might explain why the majority of stroke patients reported in recent studies are relatively cognitively preserved. In contrast, lesions to this area, in particular if they involve the MTT, would be guaranteed to lead to more pronounced cognitive outcomes.

The anterior thalamic radiations connect the anterior and mediodorsal nuclei of the thalamus with the dlPFC, vlPFC, OFC and anterior cingulate gyrus [35]. The dlPFC is associated with working memory, vlPFC with executive functions, OFC with emotional control and goal-directed behavior while the anterior cingulate gyrus is part of the Papez circuitry for mnesic functions [36]. The cortical thickness of subjects from Cluster 2 was also decreased in the left OFC, dlPFC, vlPFC and anterior cingulate gyrus; this may be related to the more frequently interrupted anterior thalamic radiations. This implies that thalamic strokes may have far-reaching implications beyond the immediate site of the lesion in some patients in accordance with the hub hypothesis.

Before concluding, however, it is worth discussing a possible limitation of our study. Nuclei supplied by more susceptible arteries, like the paramedian thalamic arteries (irrigating the mediodorsal nuclei), were overrepresented in our patient cohort, while territories served by more robust arterial systems may have been underrepresented. This bias can also be explained by the inclusion criteria from our second study including patients with lesions to the dorsomedian nuclei. It would therefore be interesting for future studies to draw insights from patients with more different arterial territories.

To conclude, our findings particularly challenge the uniformity and severity of cognitive impairments traditionally associated with thalamic lesions by demonstrating that a majority of patients had no or minor cognitive deficits. Our analysis also revealed the heterogeneity in cognitive outcomes depending on the lesion’s laterality and the specific thalamic territories impacted. Notably, lesions within the left thalamus and those affecting the mamillothalamic tract showed a significant correlation with more profound cognitive deficits, particularly in verbal memory attention and executive functions. In addition, our connectome analysis points to effects far beyond the thalamus proper of thalamic strokes. These results suggest that we should adopt a more holistic understanding of the thalamus when considering its integral role in the broader neural network governing cognitive functions.

## Data availability

The datasets generated and analyzed during the current study are not publicly available due to privacy and ethical restrictions but are available from the corresponding author upon reasonable request.

## Authors Contributions

J.PV., L.D and E.J.B designed the study.

J.P.V. performed the computations and analysis under the supervision of L.D. and E.J.B and wrote the manuscript with their support.

G.A formed J.P.V to image processing and provided technical assistance.

J.P, J.F.A, P.P, L.D, E.J.B participated to the recruitment, acquisition of neuropsychological and neuroimaging data and reviewed the manuscript.

