## Supplemental files for "Factors behind poor cognitive outcome following a thalamic stroke"

**Supplementary file**

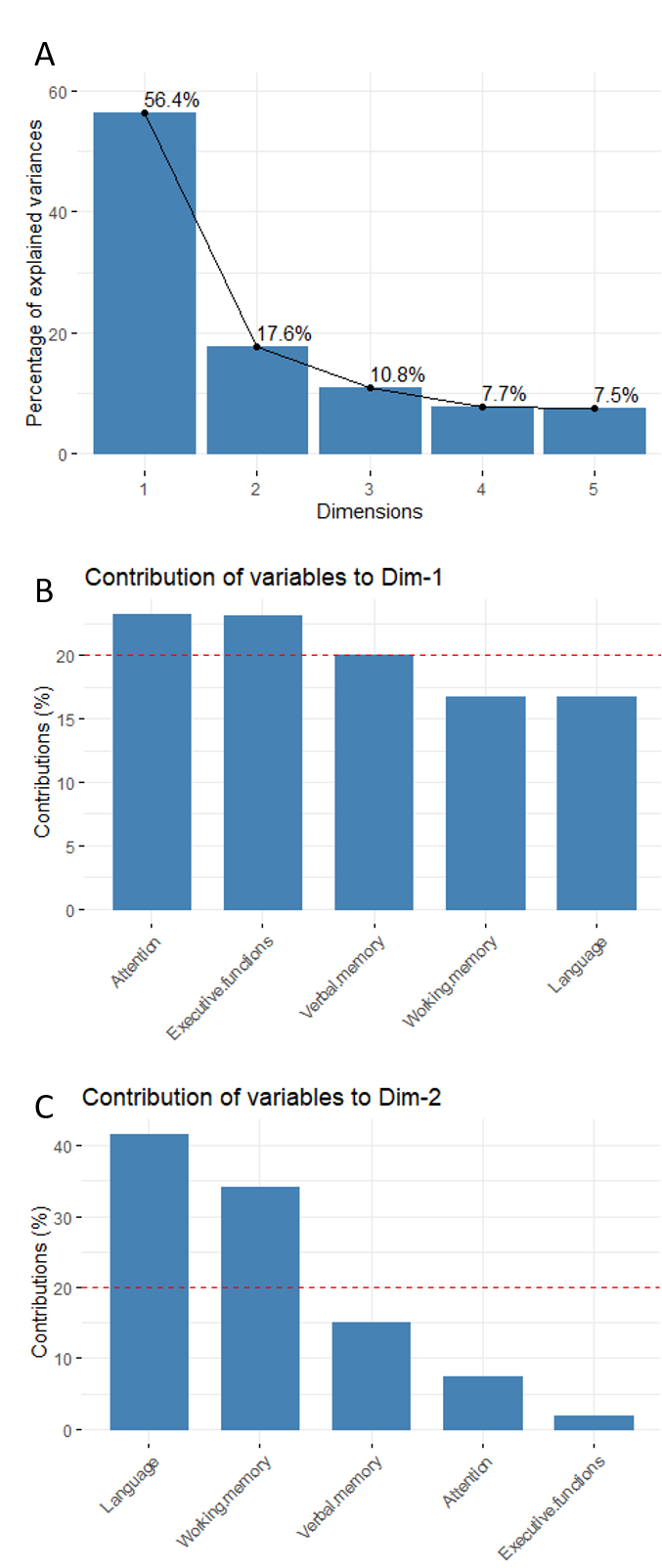

*Supplementary Figure 1: Principal component analysis. A) Percentage of variance explained of the 5 possible dimensions. B) Percentage contribution of each variable of dimension 1. C) Percentage contribution of each variable of dimension 2.*

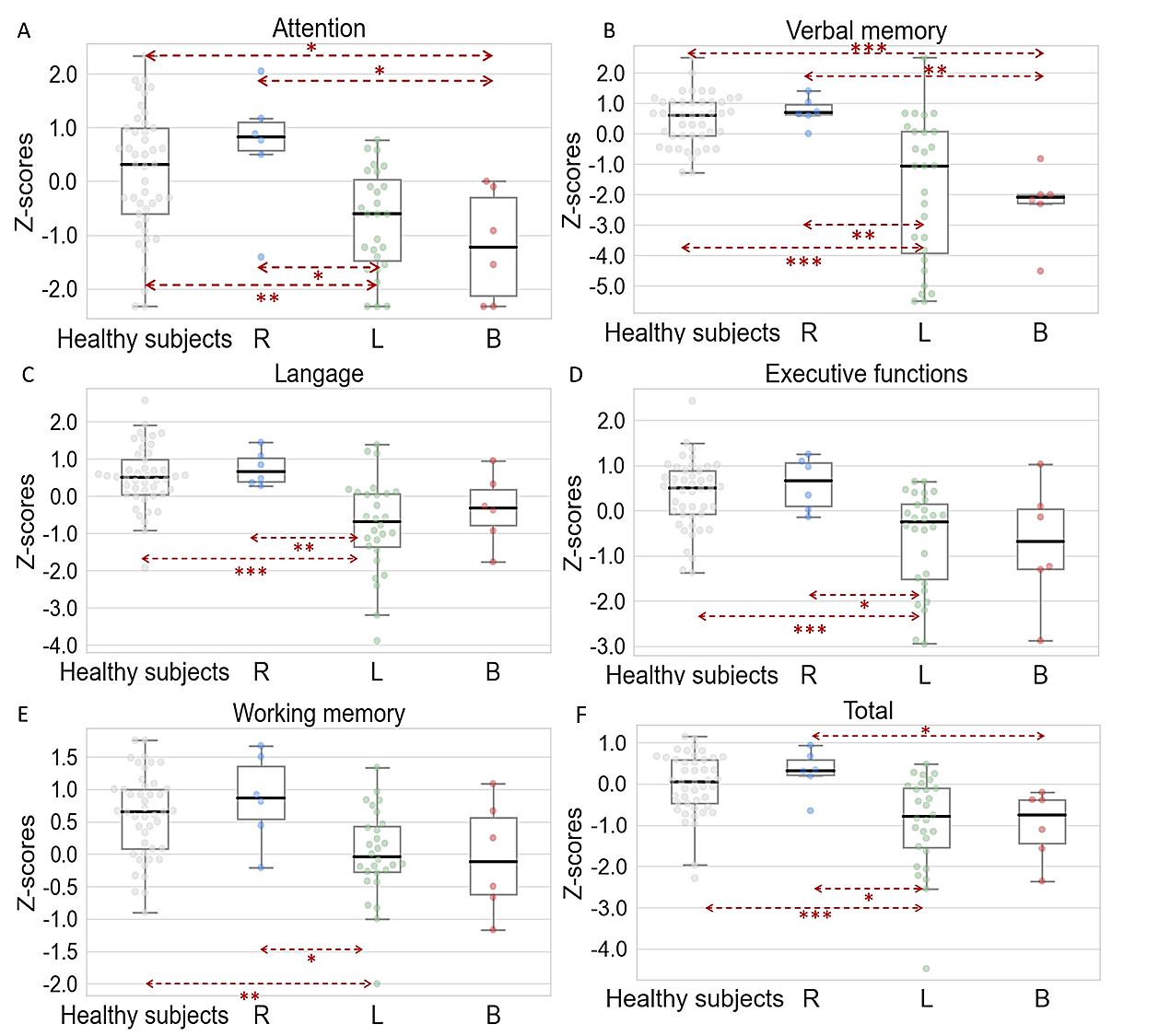

*Supplementary Figure 2: Box plot (median ± interquartile interval) of the z-scores by group of patients depending on the laterality of their infarct and healthy subjects along with a swarmplot (individual performances) of A) Attention (D2 GZ-F) B) Verbal memory (FCSRT Delayed Free Recall), C) Language (literal and semantic fluencies, confrontation naming test), D) Executive functions (digit-symbol, Stroop interference minus denomination (I-D) response time) and E) Working memory (auditory-verbal and visuospatial digit span). F) Total (mean of the previous cognitive function); R: Right (n=6), L: Left (n=28, n=27 in A), B: Bilateral (n=6); HS: Healthy Subject (n=45). Kruskal-Wallis with post-hoc Dunn’s test Bonferonni corrected for multiple comparison. * <0.05. **<0.01, ***<0.001.*
